# The impact of pH on *Clostridioides difficile* sporulation and physiology

**DOI:** 10.1101/759076

**Authors:** Daniela Wetzel, Shonna M. McBride

## Abstract

*Clostridioides difficile* is a pathogenic bacterium that infects the human colon to cause diarrheal disease. Growth of the bacterium is known to be dependent on certain bile acids, oxygen levels and nutrient availability in the intestine, but how the environmental pH can influence *C. difficile* is mostly unknown. Previous studies indicated that *C. difficile* modulates the intestinal pH, and prospective cohort studies have found a strong association between a more alkaline fecal pH and *C. difficile* infection. Based on these data we hypothesized that *C. difficile* physiology can be affected by various pH conditions. In this study, we investigated the impact of a range of pH conditions on *C. difficile* to assess potential effects on growth, sporulation, motility and toxin production in the strains 630∆*erm* and R20291. We observed pH-dependent differences in sporulation rate, spore morphology and viability. Sporulation frequency was lowest under acidic conditions, and differences in cell morphology were apparent at low pH. In alkaline environments, *C. difficile* sporulation was greater for strain 630∆*erm*, whereas R20291 produced relatively high levels of spores in a broad range of pH conditions. Rapid changes in pH during exponential growth impacted sporulation similarly among the strains. Furthermore, we observed an increase in *C. difficile* motility with increases in pH, and strain-dependent differences in toxin formation under acidic conditions. The data demonstrate that pH is an important parameter that affects *C. difficile* physiology and may reveal relevant insights into the growth and dissemination of this pathogen.

**IMPORTANCE:** *Clostridioides difficile* is an anaerobic bacterium that causes gastrointestinal disease. *C. difficile* forms dormant spores, which can survive harsh environmental conditions, allowing their spread to new hosts. In this study, we determine how intestinally relevant pH conditions impact *C. difficile* physiology in the two divergent strains, 630∆erm and R20291. Our data demonstrate that low pH conditions reduce *C. difficile* growth, sporulation, and motility. However, toxin formation and spore morphology are differentially impacted in the strains at low pH. In addition, we observed that alkaline environments reduced *C. difficile* growth, but increased cell motility. When pH was adjusted rapidly during growth, we observed similar impacts on both strains. This study provides new insights into the phenotypic diversity of *C. difficile* grown under the diverse pH conditions present in the intestinal tract, and demonstrates similarities and differences in the pH responses of different *C. difficile* isolates.

## INTRODUCTION

*Clostridioides difficile* is an emerging gastrointestinal pathogen, which often infects patients who have recently received antibiotics. Upon ingestion, the dormant spores survive the acidic pH of the stomach and enter the small intestine, where primary bile acids induce the germination of spores and enable subsequent growth of the bacterium (1–4). So far, several factors in the gastrointestinal tract are known to impact *C. difficile* growth during infection, including secondary bile acids, short-chain fatty acids (SCFA) produced by competing microbiota, host diet, host defense factors, the abundance of oxygen levels, and zinc, as well as iron and nutrient limitation (5–13). Another important factor in the gastrointestinal (GI) tract is the environmental pH, the effects of which are not well characterized for *C. difficile*. In the GI tract, the pH ranges from as low as 5.2 to as high as 7.88, depending on the region, and can be influenced by diet, transit time, health state, established microbiome and the intake of drugs (14–21). Furthermore, *C. difficile* can modulate its own environment by targeting an NHE3 ion exchanger in epithelial cells, which usually absorbs nutrients in the colon lumen by creating an H^+^-gradient (22). The loss of function of NHE3 by toxin B of *C. difficile* caused an altered intestinal environment with an increase in the luminal and fecal pH (23). A further cohort study reported a strong association between a more alkaline fecal pH and CDI, suggesting that higher pH in the GI tract may influence disease symptoms (24).

Based on these prior studies that observed *C. difficile-*directed pH changes in the intestine, we sought to investigate how the environmental pH impacts *C. difficile* physiology. To this end, we assessed the growth, sporulation efficiency, cell morphology, toxin production, motility and pH alteration *in vitro* for the historical isolate, 630Δ*erm*, and the epidemic strain, R20291. The effects of pH on growth and the ability to respond to rapid pH changes suggested a conserved mechanism for pH adaptation. However, these analyses revealed differences in the pH adaption of strains for sporulation, motility and toxin formation, which may explain differences in pathogenesis between isolates.

## MATERIALS AND METHODS

### Bacterial strains and growth conditions

Table 1 lists the bacterial strains used in this study. *C. difficile* was grown at 37°C in an anaerobic chamber (Coy Laboratory products) with an atmosphere of 10% H_2_, 5% CO_2_ and 85% N_2_ as previously described (25, 26). *C. difficile* strains were routinely cultured in brain heart infusion-supplemented (BHIS) broth or agar plates (27). To induce the germination of *C. difficile* spores, BHIS medium was supplemented with 0.1% taurocholate (Sigma-Aldrich) (28). D-fructose (0.2%) was added to overnight cultures to prevent sporulation, as needed (29).

**Table 1.**
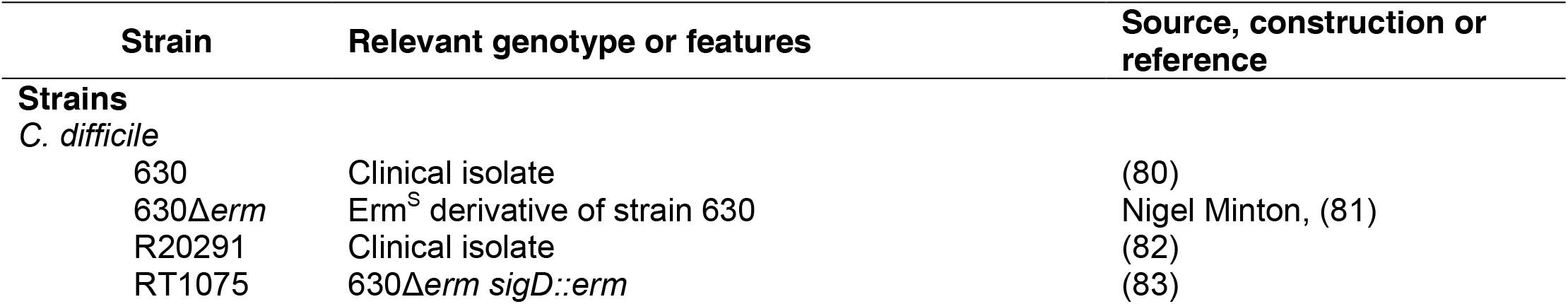
Bacterial Strains

| Strain | Relevant genotype or features | Source, construction or reference |
| --- | --- | --- |
| **Strains** |  |  |
| *C. difficile* |  |  |
| 630 | Clinical isolate | (80) |
| 630Δerm | Erm<sup>S</sup> derivative of strain 630 | Nigel Minton, (81) |
| R20291 | Clinical isolate | (82) |
| RT1075 | 630Δerm sigD::erm | (83) |

### Sporulation assay for liquid and for solid medium and phase contrast microscopy

*C. difficile* cultures were started in BHIS with 0.1% taurocholate and 0.2% fructose to induce germination of *C. difficile* spores and prevent sporulation, respectively (25, 29). To determine the sporulation efficiencies of strains in liquid cultures and on agar plates, a slightly modified 70:30 medium (29) was used without the addition of Tris base. The pH of the medium was adjusted before autoclaving using a benchtop Accumet (AB150 pH/MV meter), and the final pH was measured after complete reduction of the medium in the anaerobic chamber (30).

In brief, for sporulation in liquid medium, log-phase BHIS cultures were back-diluted in 10 ml 70:30 medium adjusted to the indicted pH for ~45 min and used to inoculate the main culture of 100 ml 70:30 medium (start OD_600_ = 0.03), which was adjusted to the same pH, respectively. The growth of strains and the pH of the culture was monitored hourly using a spectrophotometer and pH meter, respectively. At time point T2 (two hours after OD_600_ = 1.00), the total cells (vegetative cells and spores) were serially diluted and plated onto BHIS agar with 0.1% taurocholate. After 24 hours, samples were prepared for microscopy and enumeration of spores was performed as previously described (31, 32).

For sporulation on 70:30 agar plates, cultures at log-phase were diluted in BHIS broth to an OD_600_ of 0.5. These cultures (250 µl) were applied to 70:30 plates adjusted to the indicated pH, and spread as a lawn (29). After 24 hours, cells were scraped from the plates and suspended in BHIS to an OD_600_ of 1. Sample preparation, microscopy and enumeration of spores were performed as previously described (31, 32). The results represent four independent experiments and are presented as means with standard errors of the means. A one-way ANOVA and Dunnett’s test was performed for statistical comparison to the standard pH condition.

To determine the sporulation efficiency under buffered medium conditions, 70:30 medium (29) without the addition of Tris base was used. Instead of Tris, the medium was buffered using 0.1 M MES at pH 6.2, or 0.1 M HEPES at pH 7.2 or pH 8.0, respectively. The pH of the medium was adjusted before autoclaving and measured after complete reduction, as described for broth medium. Cultures of *C. difficile* grown in BHIS broth were back-diluted into 10 ml 70:30 medium, which was adjusted to the indicated pH, and used to inoculate the main culture of 100 ml 70:30 medium (OD_600_ = 0.03) at the same pH, respectively. At time point T2 as described above for the liquid medium, total cells were diluted, plated onto BHIS agar with 0.1% taurocholate, and enumerated.

### Sporulation assay under rapid changing pH conditions

To determine sporulation frequencies under changing pH conditions, 70:30 medium without the addition of Tris base was used. For liquid medium, log-phase growing *C. difficile* in BHIS were back-diluted into 10 ml of 70:30 medium adjusted to a pH of 7.2, which was used to inoculate the main culture of 100 ml 70:30 medium at pH 7.2 (start OD_600_ = 0.03). The growth of strains and pH of the culture was monitored hourly. At an OD_600_ of 0.5, the pH (~pH 6.8) in the cultures was rapidly changed by the addition of 5 N HCl or 5 N NaOH to obtain an increase or decrease of 0.5 U (units), 1.0 U and 1.5 U in pH, respectively. At time point T2, as described above for liquid medium, total cells were diluted and plated onto BHIS agar with 0.1% taurocholate. Sample preparation, microscopy, and enumeration of spores were performed as described above.

To assess sporulation efficiencies under rapid pH changes on 70:30 plates, log-phase cultures of *C. difficile* were back-diluted into BHIS to an OD_600_ of 0.5 and 250 µl of sample were applied to 70:30 plates at a pH of 6.8 by spreading them as a lawn. After six hours of growth, the cells were scraped from the plate into BHIS, adjusted to an OD_600_ of 0.5, and 250 µl were applied to 70:30 plates at a pH of 6.8 as an internal control, and to 70:30 plates with 0.5 U, 1.0 U, and 1.5 U increases in pH, or 0.5 U, 1.0 U, and 1.5 U decreases in pH, respectively. After 24 hours, samples were prepared for microscopy and enumeration of spores was performed as described above. The results represent four independent experiments and are presented as means with standard errors of the means. A one-way ANOVA and Dunnett’s test was performed for statistical comparison to the standard pH condition.

### Swimming motility Assay

Strains were grown overnight in BHIS with 0.1% taurocholate and 0.2% fructose. Active cultures were diluted and grown to an OD_600_ of 0.5 in BHIS broth, and 5 µl of culture was stabbed in the center of one-half concentration BHI plates with 0.3% agar (33). The pH of the BHI plates was adjusted and measured as described above. The cell growth in diameter was measured every 24 h for five days. The results represent four independent experiments and are presented as means with standard errors of the means. A one-way ANOVA and Dunnett’s test was performed for statistical comparison to the standard condition of pH 7.2.

### Western blot analysis

Strains were grown in BHIS broth to an OD_600_ of 0.5 and 250 µl of cultures were applied as a lawn to 70:30 plates at different pHs, as indicated. After 24 h of incubation, cells were scraped from the plates, suspended in SDS-PAGE loading buffer, and processed as previously described (33, 34). Total protein was quantified using the Pierce Micro BCA Protein Assay Kit (Thermo Scientific). 8 µg of total protein was separated by electrophoresis on a precast 4-15% TGX gradient gel (Bio-Rad) and transferred to a 0.45 µm nitrocellulose membrane. Western blot analysis was conducted using a mouse anti-TcdA antibody (Novus Biologicals), followed by a goat anti-mouse Alexa Fluor 488 (Life Technologies) secondary antibody. Imaging and densitometry were performed with a ChemiDoc and Image Lab Software (Bio-Rad). A one-way ANOVA, followed by Dunnett’s multiple-comparison test, was performed to assess statistical differences in TcdA protein levels compared to the standard condition of pH 7.0. Four independent biological replicates were analyzed for each strain.

## RESULTS

### pH impacts *C. difficile* growth and spore formation differentially in 630∆*erm* and R20291

To assess the impact of the environmental pH on *C. difficile*, we examined the genetically distinct strains 630Δ*erm* and R20291 in 70:30 sporulation broth in a range of pH that is physiologically relevant to the large intestine. Cultures were monitored for effects on growth, change in pH overtime, and sporulation in medium at pH 6.2, 7.2, and 8.0, respectively (Fig. 1). For both strains, significant decreases in growth were observed during mid-logarithmic growth at the acidic pH 6.2 and at the alkaline pH 8.0, relative to pH 7.2 (Fig. 1A, B). Analyzing the change in pH of the cultures, the largest drop in pH could be seen for pH 8.0 and pH 7.2 cultures, which decreased from pH 8.0 to 7.4, and from pH 7.2 to 6.5, within 8 hours for both strains. For pH 6.2 cultures, similar decreases were observed for both strains during growth, with decreases from pH 6.2 to 5.8 within 6 hours, respectively (Fig. 1A, B). In addition, in the pH 7.2 and 6.2 cultures, the pH increased after ~6 hours, around the transition to stationary phase growth.

**Figure 1.**
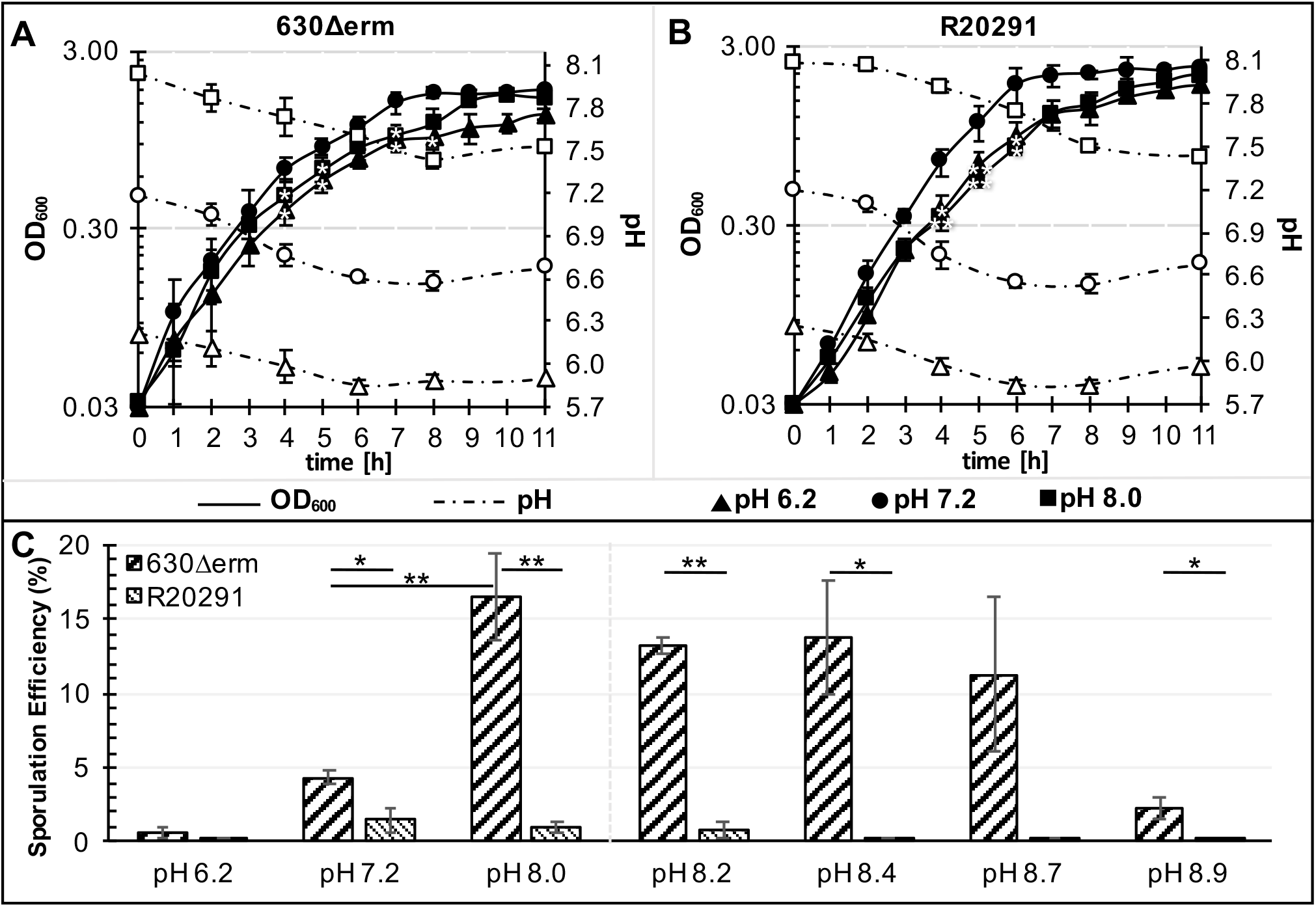
Growth and sporulation of *C. difficile* is impacted by pH in liquid sporulation medium. The strains of 630Δ*erm* (A) and R20291 (B) were cultivated in 70:30 broth at a pH of 6.2, pH 7.2, or pH 8.0. (C) Analysis of the sporulation efficiency of strains grown in pH 6.2, 7.2, and pH 8.0, or in high alkaline pH cultures of 8.2, 8.4, 8.7, and pH 8.9, respectively. Data were analyzed by Student’s two-tailed *t*-test comparing strains of 630∆*erm* and R20291, and by one-way ANOVA with Dunnett’s test for multiple comparisons to pH 7.2. * indicates P value of ≤ 0.05; ** indicates adjusted P value of ≤ 0.01; n=4.

Analysis of the spore formation under the different pH conditions uncovered strain-dependent differences in sporulation efficiency (Fig. 1C). The 630Δ*erm* strain produced more spores in sporulation broth than the R20291 strain under every pH condition tested. Additionally, 630Δ*erm* demonstrated dramatic increases in sporulation frequency as the pH increased (pH 8.0, ~17% vs ~4% at pH 7.2 and less than 1% at pH 6.2) (Fig. 1C). In comparison, R20291 demonstrated relatively low sporulation under all pH conditions (Fig. 1C). Both strains exhibited very low spore formation at pH 6.2, suggesting that acidic conditions do not support sporulation in *C. difficile* (~0.7% for 630Δ*erm* and ~0.1% for R20291).

Based on the correlation observed between spore formation and higher pH, we further assessed growth and sporulation under more alkaline pHs (pH 8.0 to 8.9). Despite similar impacted growth of strains under high alkaline pH conditions (Fig. S1A, B), R20291 sporulated at less than 1% efficiency, with very few spores formed above pH 8.2 (Fig. 1C). In comparison, 630Δ*erm* maintained robust spore formation up to pH 8.7 and only reduced sporulation at a pH of 8.9 to ~2% (Fig. 1C).

To investigate if the initial pH of the medium or bacterial-dependent changes in pH impact sporulation, we limited the change in pH by buffering the culture medium. Buffers appropriate for the respective pH conditions were utilized and sporulation assessed as described in Material and Methods (0.1 M MES at a pH of 6.2, or 0.1 M HEPES at pH 7.2 and pH 8.0). As expected, buffering the medium limited the pH shift of cultures over time; however, it did not alter the growth or sporulation of either strain (**Fig. S2**). These data suggest that although growth of the strains appears similar in different pH conditions, the effects of pH on sporulation are considerably greater for the 630Δ*erm* strain than for R20291.

Because of the observed pH effects on *C. difficile* growth and sporulation in liquid cultures, we asked how pH impacts sporulation on solid medium, which is known to support more robust spore formation (32, 34–36). To test this, sporulation agar plates were prepared with a range of pH (pH 5.2 to 9.0), and the spore formation for 630Δ*erm* and R20291 was determined. Both strains produced considerably more spores on solid medium than were observed in liquid (Fig. 1, Fig. 2). Consistent with results from broth cultures, the strains showed differences in spore production under different pH conditions. At a low pH of 5.2, both strains were inhibited for growth (Fig. 2). Growth of both strains was observed at a pH of 5.5; however, this low pH resulted in a relatively low sporulation efficiency of 8-9% for both strains, compared to the 31-34% sporulation observed under neutral pH (pH 7.0, Fig. 2). The strain 630Δ*erm* exhibited increased sporulation with increases in pH, reaching a maximal sporulation efficiency at pH 8.0 of ~38% (Fig. 2). In comparison, R20291 demonstrated maximal sporulation efficiency at a pH of 6.0 and exhibited a broader pH range for robust spore formation (pH 6.0 to pH 8.5, Fig. 2). Beyond pH 8.0, both strains exhibited a stepwise reduction in spore formation. Overall, the data suggest that in liquid or solid medium, 630Δ*erm* sporulated best in more alkaline pH, whereas R20291 sporulated robustly across a broad range of pH conditions.

**Figure 2.**
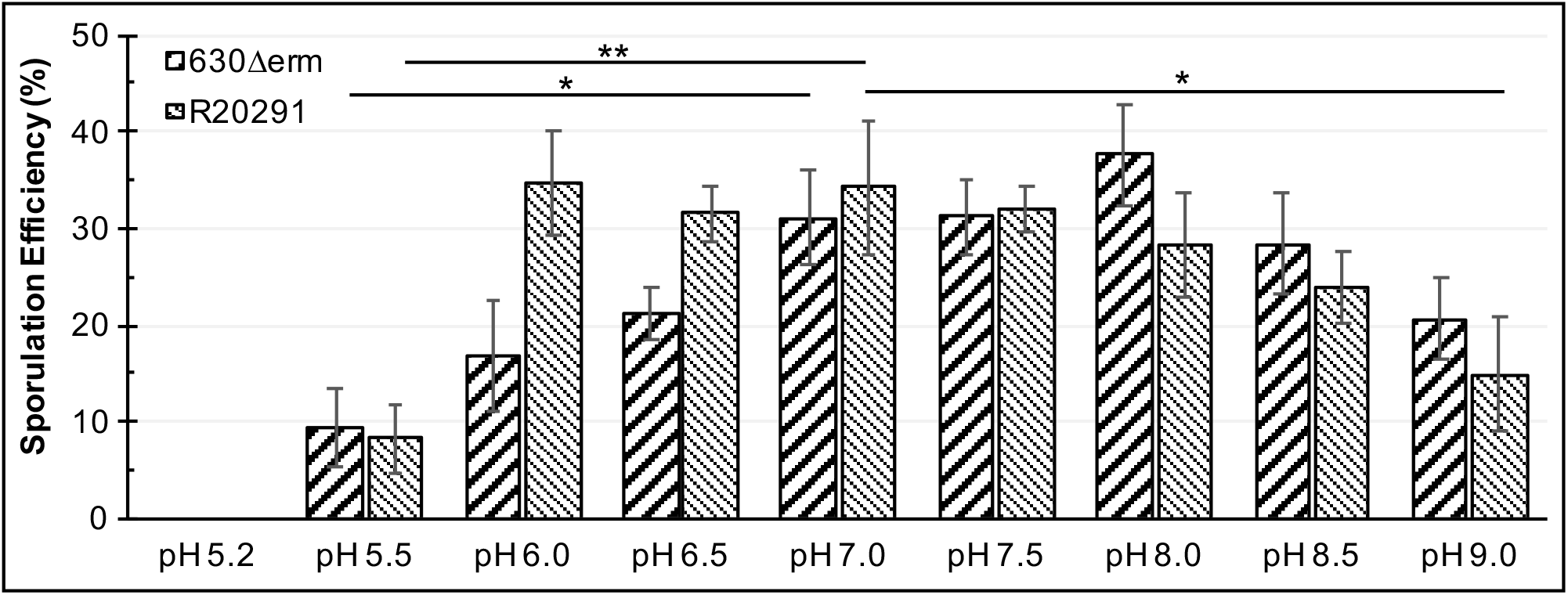
Sporulation of *C. difficile* on solid medium is influenced by pH. Strains 630Δ*erm* and R20291 were analyzed for sporulation on 70:30 plates at the indicated pH. No growth or sporulation was detected at pH 5.2. Data were analyzed by one-way ANOVA with Dunnett’s test for multiple comparisons to pH 7.0, indicated by brackets. * indicates P value of ≤ 0.05; ** indicates P value of ≤ 0.01; n=4.

To assess the effects of pH on vegetative cell and spore morphology, phase-contrast microscopy was performed for strains grown in different pH conditions (Fig. 3). Both strains exhibited slightly elongated vegetative cells and the lowest phase-bright spore formation under low pH conditions. Additionally, R20291 formed small, round, phase-dark spores and exhibited more pronounced lysis of cells at pH 6.2 in liquid, compared to the other pH conditions. These results suggest that R20291 cells initiate sporulation at pH 6.2 in liquid, but form few mature, ethanol-resistant spores under these conditions. However, R20291 does not produce these small, immature spores on solid media at low pH, suggesting that the viscosity of the medium plays a role in spore formation. 630Δ*erm* produced the most abundant phase-bright spores at pH 8.0 for both liquid and solid media, consistent with the results of ethanol-resistance spore assays (Fig. 1, Fig. 2).

**Figure 3.**
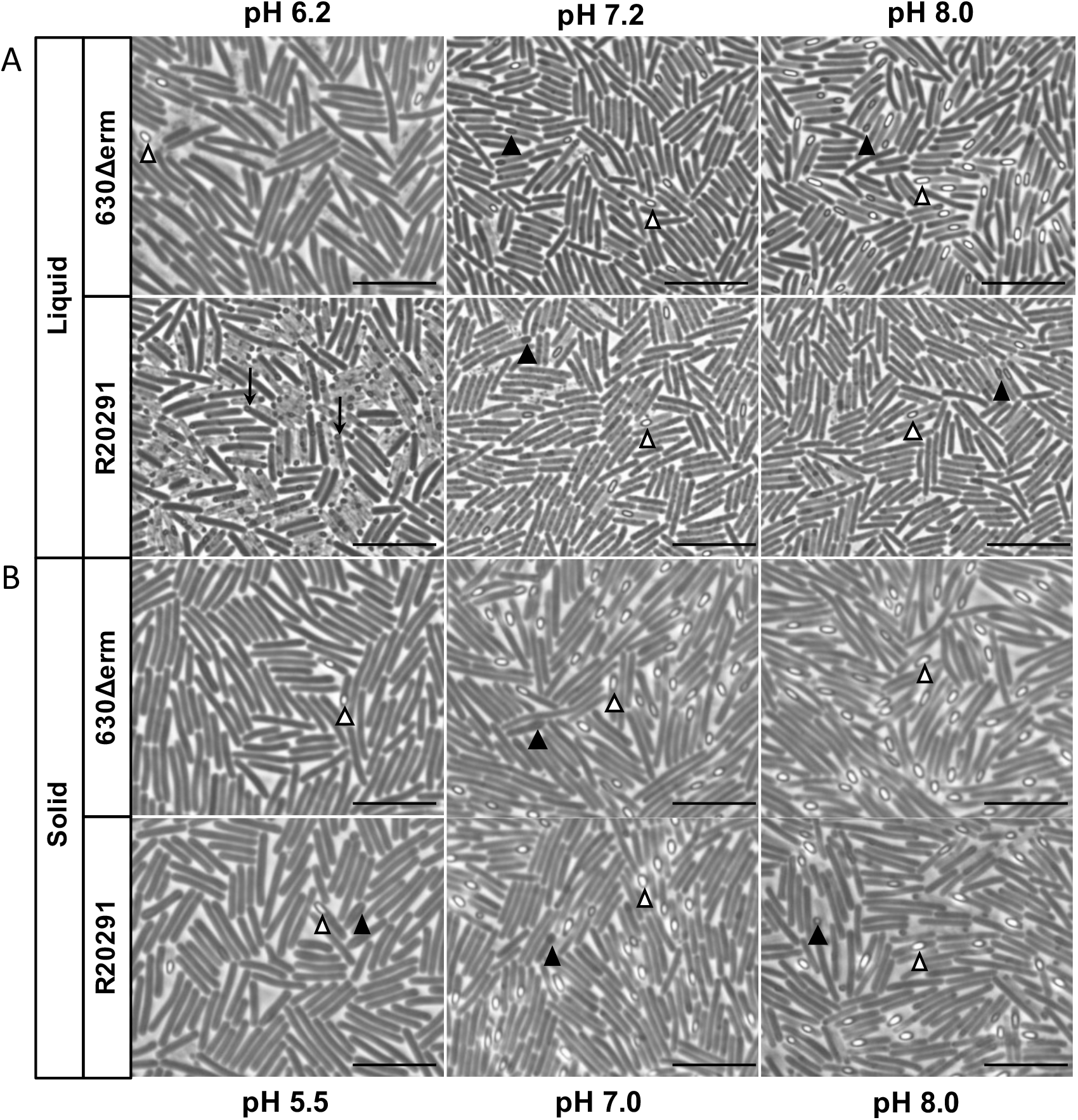
Phase-contrast microscopy of *C. difficile* strains grown under diverse pH conditions revealed differences in spore morphology and abundance. In liquid culture (A) and on solid medium (B) after 24 h of growth, 630∆*erm* and R20291 were analyzed for cell morphology. Filled arrowheads (▲), phase-dark pre-spores; open arrowheads (∆), phase-bright mature spores; arrow (**↓**), small immature spores. Bars = 10 µm.

### Rapid changes in pH impact growth and sporulation for both strains and are dependent on the medium viscosity

As the pH varies greatly throughout large intestine, *C. difficile* is exposed to abrupt shifts in pH during transit through the colon. Considering this, we investigated the impact of rapid changes on growth and spore formation. *C. difficile* was cultivated at a moderate pH in broth or on agar medium (Fig. 4), and exponentially growing cells were exposed to rapid increases or rapid decreases in pH. For the pH shift experiment in liquid medium, 630Δ*erm* and R20291 cultures were grown to an OD_600_ of 0.5 (pH ~6.8), then the pH of the culture was shifted by the addition of HCl to decrease the pH 0.5, 1.0, and 1.5 units, or by the addition of NaOH to increase the pH 0.5, 1.0, and 1.5 units, respectively (**Fig. S3**). Growth of both strains was minimally impacted by pH shifts of +/− 0.5 units (**Fig. S3**). A moderate reduction in growth was observed for both strains when the pH increased 1.0 or 1.5 units. In comparison, a decrease of 1.0 pH unit drastically impacted growth of both strains, and a 1.5 unit decrease resulted in extinction of the growth (**Fig. S3**). After a decrease of 1.0 pH unit, growth was inhibited for the next three hours and the culture densities were significantly lower than the unadjusted culture (**Fig. S3A, B**). Sporulation was evaluated in pH-adjusted cultures after 24 h of growth and all changes in pH decreased spore formation in both strains (Fig. 4A). As expected, the decreases in sporulation efficiencies correlated directly with the magnitude of the pH shift. These results indicate that *C. difficile* is able to survive relative changes in the pH of an aqueous environment, but with significant decreases in spore production.

**Figure 4.**
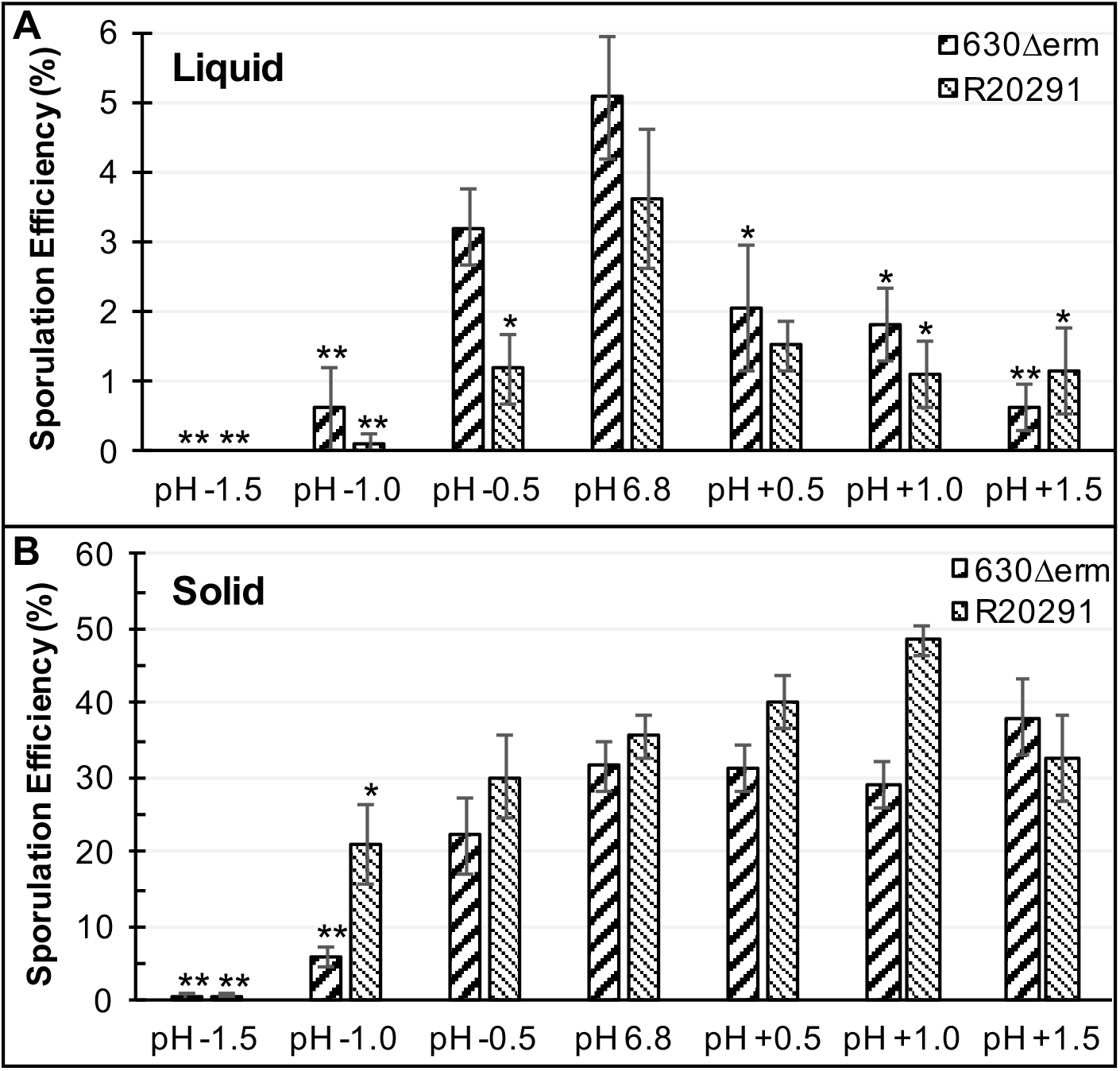
A rapid change in pH influences sporulation of *C. difficile* strains. (**A**) Strains 630Δ*erm* and R20291 were cultivated in 70:30 broth medium at an initial pH of 7.2 and upon reaching an OD_600_ of ~0.5, the pH (~6.8) was rapidly shifted (+/−) 0.5 U, 1.0 U, and 1.5 U, as indicated. Sporulation efficiency was assessed after 24 h of growth. No survival of cells or sporulation was detected when the pH was rapidly decreased 1.5 U for either strain in liquid. n=4. (**B**) After 6 h of growth on sporulation agar, cells of 630Δ*erm* and R20291 were transferred to sporulation agar at pH 6.8 (control) or to plates with pH shifted (+/−) 0.5, 1.0, or 1.5 units. Sporulation was determined after 24 h of growth. Data were analyzed by one-way ANOVA and Dunnett’s test for multiple comparisons to the starting pH. * indicates P value of ≤ 0.05; ** indicates adjusted P value of ≤ 0.01; n=4.

To test if rapid changes in pH can impact sporulation during growth on solid surfaces*, C. difficile* strains were similarly cultivated on 70:30 plates at a pH of 6.8 for six hours, then transferred to a second pH 6.8 plate, as an internal control, or to 70:30 plates with 0.5, 1.0, and 1.5 unit increases or decreases in pH, respectively (Fig. 4B). In contrast to growth in liquid medium, sporulation of both strains was marginally impacted when the pH was increased by 0.5, 1.0, or 1.5 units. Conversely, decreases of 1.0 or more pH units impacted sporulation of either strain, with the greatest decrease in spores observed for the 630Δ*erm* strain. The data demonstrate that the higher viscosity of solid medium enhances *C. difficile* adaptation to abrupt pH shifts. Overall, *C. difficile* strains adapted better to increases in pH than to decreases in pH, regardless of the viscosity of the medium, as evidenced by growth and sporulation under all increased pH conditions. In comparison, decreases in pH drastically impaired *C. difficile* growth and spore formation (Fig. 4A, B). Both strains were able to decrease the pH of the culture when the pH was increased, lowering the pH to a more neutral value (**Fig. S3C, D**). In contrast, when the pH was rapidly decreased, neither strain could effectively increase the pH over time, resulting in prolonged and deleterious exposure of the cells to acidic conditions.

### Toxin formation is impacted by pH in a strain-dependent manner

Since sporulation and toxin expression are co-regulated in *C. difficile* (33, 34, 36–39), we considered that the pH during growth may also influence toxin production. Thus, we assessed the production of toxin A from strains grown on 70:30 plates in a range of pH conditions (pH 5.5 to 8.5) by western blot (Fig. 5). Under acidic conditions, the 630Δ*erm* strain exhibited reduced toxin, relative to production at pH 7.0. In contrast, the R20291 strain demonstrated increased toxin A formation under acidic conditions (Fig. 5). Toxin production for 630Δ*erm* was greatest at pH 6.5 and 7.0, while R20291 produced the most toxin at a pH range of 5.5 to 7.0. Both strains exhibited lower toxin production under alkaline conditions, which is in contrast to spore production in the same conditions (Fig. 4B, Fig. 5). These data suggest that sporulation and toxin expression are differentially controlled under alkaline conditions.

**Figure 5.**
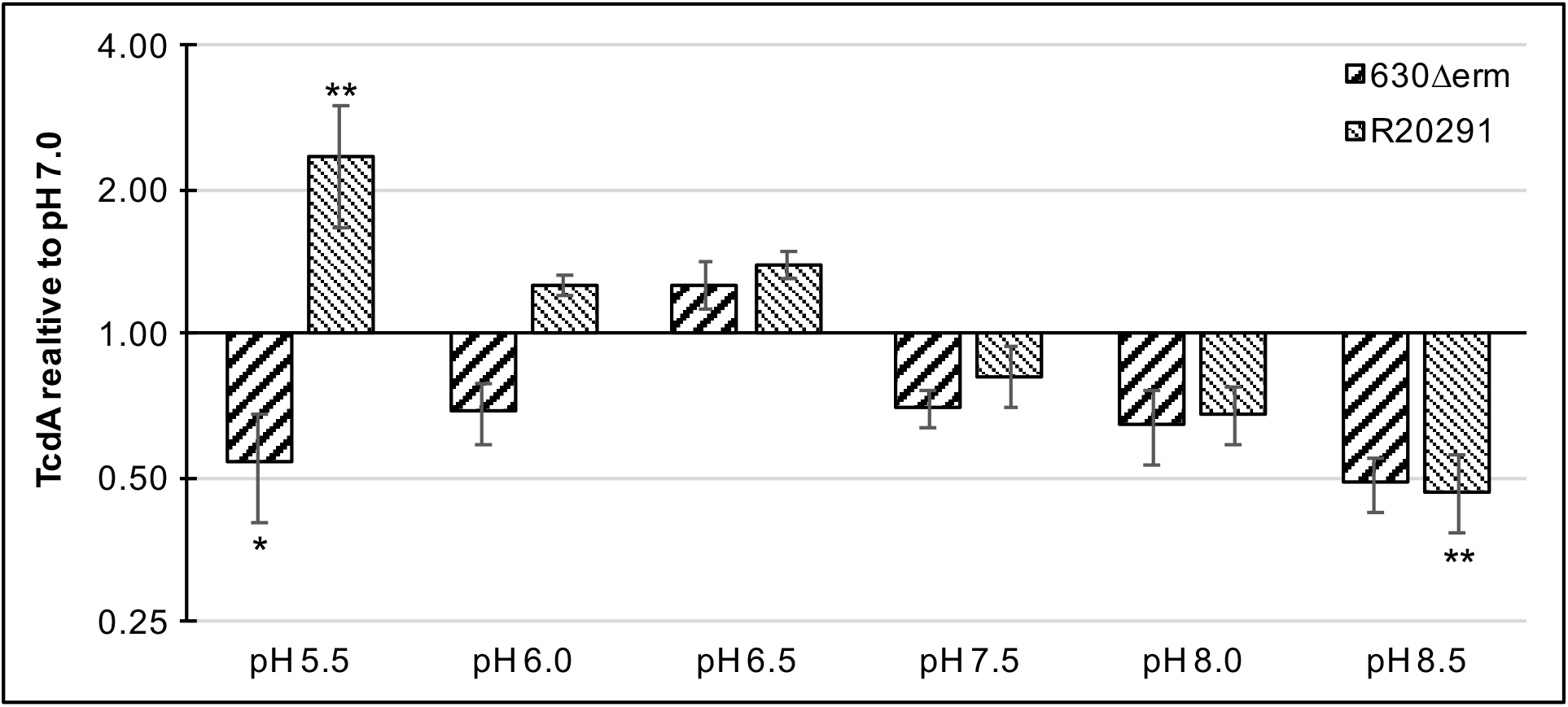
Strain-dependent differences in toxin A production under different pH conditions. Strains 630Δ*erm* and R20291 were analyzed for TcdA production on 70:30 plates at the indicated pH by western blot analysis after 24 h of growth. Densitometries were normalized to pH 7.0; Scale plotted at log2. Analyzed by one-way ANOVA and Dunnett’s test compared to growth at pH 7.0. * indicates P value of ≤ 0.05; ** indicates P value of ≤ 0.01; n=4.

### *C. difficile* motility is affected by pH

Because toxin expression and motility are both regulated by SigD, the flagellar alternative sigma factor (40–42), we next investigated the impact of pH on *C. difficile* motility. To test this, we analyzed the motility of *C. difficile* on soft agar plates at a range of pH conditions between 6.2 and 8.2. Swimming motility was measured every 24 hours for five days, and the non-motile *sigD* mutant strain (RT1075) was used as a negative control (Fig. 6). Both strains demonstrated the least motility at low pH (6.2) and the greatest motility at pH 7.7-8.2 (Fig. 6A, B). However, strain-dependent differences in the distance travelled were observed, with 630Δ*erm* exhibiting greater motility than R20291 at pH 6.2 and 6.7, and R20291 surpassing 630Δ*erm* at pH 7.2, 7.7, and 8.2. The poor motility for the R20291 strain at low pH contrasts with the higher toxin production observed at low pH for this strain, suggesting that in R20291, factors other than SigD activity restrict motility under acidic conditions. These data demonstrate that pH is an important environmental factor that influences *C. difficile* motility, and that the bacteria are generally more motile in alkaline conditions and less motile at lower pH.

**Figure 6.**
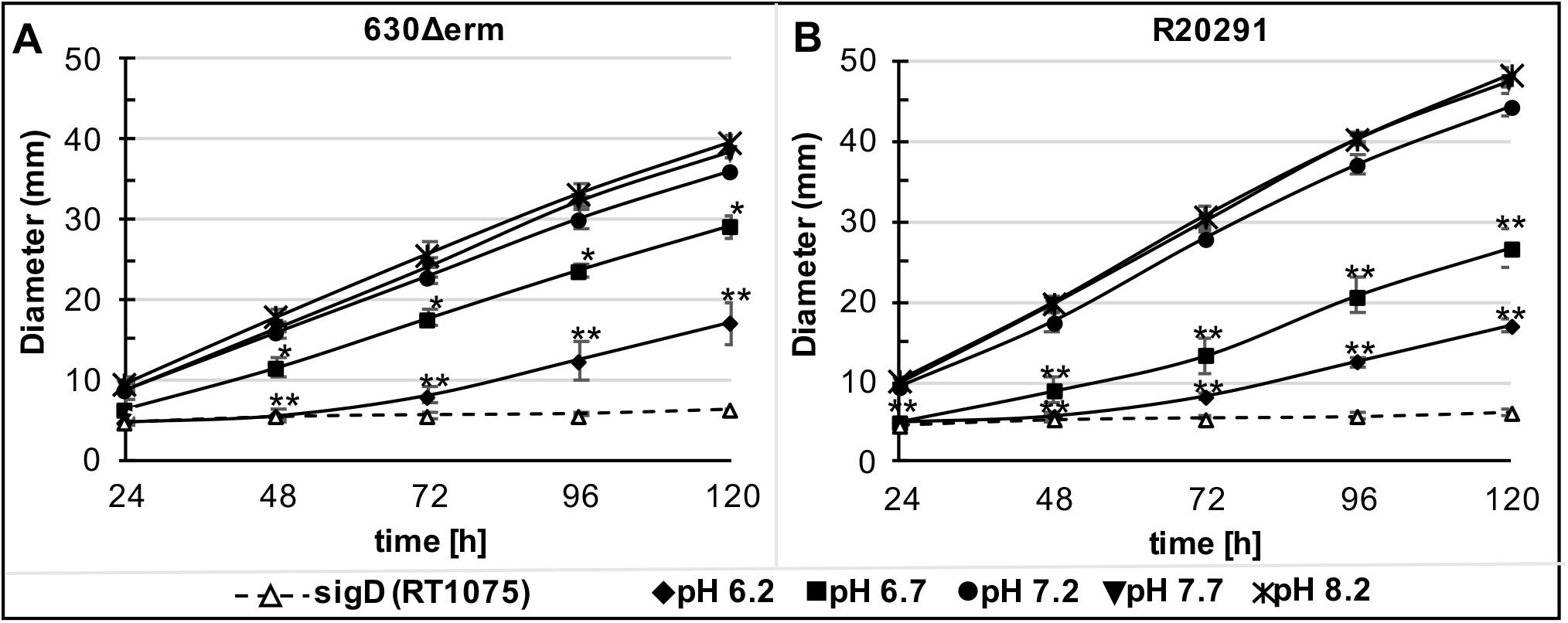
*C. difficile* motility is impacted by pH condition. Strains 630Δ*erm* (A), R20291 (B), and a *sigD* mutant (RT1075, negative control) were tested for motility on ½ BHI plates (0.3% agar) at pH 6.2, 6.7, 7.2, 7.7, and pH 8.2, respectively, by measuring the swimming motility every 24 h for 120 h. Significantly reduced motility was observed at pH 6.2 and pH 6.7, relative to their respective motility at pH 7.2, for both strains. Data were analyzed by one-way ANOVA and Dunnett’s test compared to pH 7.2. * P ≤ 0.05; ** adjusted P value of ≤ 0.01; n=4.

## DISCUSSION

*C. difficile* is exposed to diverse and changing pH conditions during transit through the gastrointestinal tract. Prior studies showed that the environmental pH impacts spore germination, resulting in inhibition of germination under acidic conditions that is reversed upon exposure to neutral pH (1, 30, 43, 44), which highlights a limiting factor for the germination of spores within the small bowel in the presence of bile acids. In this study, we further assessed the impact of diverse pH conditions on *C. difficile* growth, sporulation, toxin formation, and motility.

As strain-dependent phenotypic differences are often described in *C. difficile* (34, 36, 37, 42, 45–47), we investigated the *C. difficile* pH response using the historical isolate 630∆*erm*, which is commonly studied in laboratories, and the epidemic-associated strain R20291. We found that during growth in liquid culture, no strain-dependent differences were observed under different pH conditions (Fig. 1A). However, spore formation differed between the strains grown in broth culture, as evidenced by a dramatic increase in spore production for strain 630∆*erm* in alkaline conditions (Fig. 1B, C), and only modest sporulation for R20291 at any pH. Thus, despite the greater culture density achieved by R20291 in liquid medium (Fig. 1), sporulation in this strain is hindered in broth culture. Differences in pH-dependent sporulation were also seen on solid sporulation medium, as 630∆*erm* showed maximal sporulation from pH 7.0 to 8.5, while R20291 sporulated best at a range of pH from 6.0 - 8.5.

Under acidic conditions in liquid and on solid medium, both strains produced significantly fewer mature spores. Decreases in spore formation in acidic conditions has also been described for *Bacillus* spp., including *B. weihenstephanensis* at pH 5.6, *B. licheniformis* at pH 6.3, and *B. cereus* at pH 5.9-6.1 (48–52). The clostridia appear more varied in their sporulation efficiency response to pH. In *C. perfringens*, spores are produced within a narrow pH range of 5.9 to 6.6 (53). In contrast, *C. cellulolyticum* reached highest sporulation efficiency at the lowest tested pH of 6.4 (54). In accordance with previous studies, no growth of *C. difficile* was observed below a pH of 5.5 (55–58). Comparative genomic analysis of *C. difficile* versus *C. sordellii* revealed the absence of several acid adaptation and broad range pH survival mechanism in *C. difficile*, such as ureases, glutamate decarboxylase, arginine deaminase or potassium transport proteins (59), which may explain the inability of *C. difficile* to grow under more acidic conditions.

Usually *C. difficile* infects the colon, but cases of *C. difficile* enteritis involving the small bowel have been described (60, 61) (62, 63). From the proximal small bowel to the ascending colon the pH drops dramatically from 7.88 to 5.26, due to fermentation from colonic bacteria and the production of short-chain fatty acids (64–66). Between the ascending colon and the descending colon the pH gradually increases as a result of mucosal bicarbonate secretion, colonic ion channel activity and Na^+^/H^+^ exchange (23, 67, 68). To understand how these pH fluctuations impact *C. difficile*, we investigated the vegetative cell responses to rapid changes in pH. We found that rapid pH changes similarly impacted the growth and spore formation of both strains (**Fig. S3**, Fig. 4). In general, both strains adapted better to rapid increases in pH than to pH decreases, which could be explained by deficiencies in acid adaptation mechanisms (59). Emerson et al., 2008 analyzed the transcriptional response of *C. difficile* strain 630 after acid or alkaline shocks of 1.5 pH units. Acid shock induced genes of the general stress response, as well as the heat shock regulon of HrcA, CtsR, GroEL and GroES (69), which indicates a severe stress for *C. difficile*. However, the authors did not test if *C. difficile* could survive the acid or alkaline shock treatments (69). In this study, we found that *C. difficile* will adapt and sporulate on agar medium after an alkaline shock of 1.5 units, but cannot survive an acid shock of 1.5 units in broth cultures. Although both strains survived a 1.5 unit acid shock on solid medium, it resulted in drastic reduction in spore formation (Fig. 4).

Differences in phenotypes for cells grown on solid and liquid medium have been observed for *B. cereus*, with plate-grown cells displaying increased gamma radiation resistance and a more developed S-layer compared to cells grown in liquid (70). Other studies of *C. difficile* found substantial differences in gene expression profiles between biofilms grown in broth and grown on plates (71), as well as differences in the expression of phase variation genes and the orientation of invertible elements (42, 72). We anticipate that differences in gene expression for liquid and solid medium affected the survival and adaptation of *C. difficile* after an acid shock; however, this was not explored in this study.

Previous studies that examined the effect of pH on *C. difficile* toxin formation also noted strain-dependent differences in toxin production. For the *C. difficile* VPI 10463 strain (ribotype 087), toxin formation after 24 h and 48 h was highest for cells grown at pH 6.5-7.5, and reduced at pH 5.5 and 8.5, respectively (56). In our study, we analyzed the toxin formation after 24 h of growth on 70:30 sporulation agar across a range of pH, and found strain-dependent differences in toxin formation at different pH. Like the VPI 10463 strain, strain 630∆*erm* produced the most toxin at a pH of 6.5, and significantly reduced toxin production at pH 5.5 and 8.5. R20291, in comparison, significantly increased toxin formation at a pH of 5.5, and demonstrated the greatest toxin formation at a pH range of 5.5-7.0. The increased toxin formation of R20291 (027 ribotype) at a low pH 5.5 is in accordance with prior observed increase in the expression of toxin genes in an clinical isolate of 027 grown at pH 5 (73).

Our data indicate that low pH conditions reduced motility and high pH increased motility for *C. difficile* (Fig. 6), similar to observations in *B. subtilis* (74). As described previously (75), a possible explanation for the reduced motility under acidic conditions, could be the necessity to close the flagella motor-driven proton entrance and energy costs for flagellum biosynthesis under harmful conditions (76, 77). Another possible explanation for reduced motility may be the incorrect assembly of flagellar proteins or protein instabilities under low pH conditions, as described elsewhere (78, 79).

In this study, we identified pH-dependent effects on strain growth, toxin production, motility, and sporulation, all of which can be used to improve growth and phenotypic testing in diverse *C. difficile* isolates. We discovered differences in the pH adaptation of strains, and impacts of the medium viscosity on *C. difficile.* This ability of R20291 to adapt to a broad range of pH conditions for sporulation and toxin production (Fig. 2, Fig. 5), may provide the strain with an advantage for host colonization and pathogenesis. In strain 630∆*erm*, motility and toxin formation were similarly depressed in low pH conditions. This is in contrast to R20291, which demonstrated the greatest toxin formation at low pH, but less motility under low pH conditions. Further studies are necessary to understand how *C. difficile* strains regulate these individual processes in response to pH changes within the host.

## ACKNOWLEDGEMENTS

We give thanks to members of McBride lab for helpful suggestions and discussions during the course of this work. This research was supported by the U.S. National Institutes of Health through research grants AI116933 and AI121684 to S.M.M. The content of this manuscript is solely the responsibility of the authors and does not necessarily reflect the official views of the National Institutes of Health.

---

**Figure S1. Growth of *C. difficile* is impacted with increasing alkaline pH in liquid culture.** Strains 630∆*erm* (A) and R20291 (B) were cultivated in liquid 70:30 broth at a pH of 8.0, 8.2, 8.4, 8.7, and pH 8.9, respectively. Data were analyzed by one-way ANOVA and Dunnett’s test compared to pH 8.0. * indicates P value of ≤ 0.05; ** indicates adjusted P value of ≤ 0.01; n=3.

**Figure S2. Growth and sporulation of *C. difficile* is similarly impacted by pH under buffered medium conditions.** 630Δ*erm* (A) and R20291 (B) were cultivated in 70:30 cultures at a pH of 6.2 with or without 0.1 M MES, or at pH 7.2 or 8.0 with or without 0.1 M HEPES, respectively. Only for R20291, a pronounced decrease in growth in buffered pH 8.0 culture compared to the unbuffered pH 8 culture was observed (* P ≤ 0.05; ** P ≤ 0.01). (C) Sporulation of strains under buffered medium conditions compared to non-buffered cultures. No statistical differences were observed. Data were analyzed by Student’s two-tailed *t*-test. Experiments were performed three or more times.

**Figure S3. A rapid change in pH influenced growth of *C. difficile* in liquid culture.** Strains 630Δ*erm* (A,C) and R20291 (B,D) were cultivated in 70:30 broth at an initial pH of 7.2. At an OD_600_ of 0.5 (indicated by arrow), the pH was increased or decreased 0.5, 1.0, or 1.5 pH units, respectively. A significant decrease in growth could be observed when the pH was rapidly shifted (+/−) 1.0 U or −1.5 U. Data were analyzed by one-way ANOVA with Dunnett’s test for multiple comparisons to growth at pH 7.2. * indicates P value of ≤ 0.05; ** indicates adjusted P value of ≤ 0.01, n=3.

